# AptaMat: a matrix-based algorithm to compare single-stranded oligonucleotides secondary structures

**DOI:** 10.1101/2022.05.04.490414

**Authors:** Thomas Binet, Bérangère Avalle, Miraine Dávila Felipe, Irene Maffucci

## Abstract

Comparing single-stranded nucleic acids (ssNAs) secondary structures is fundamental when investigating their function and evolution and predicting the effect of mutations on the ssNAs structures. Many comparison metrics exist, although they are either too elaborate or not enough sensitive to distinguish close ssNAs structures.

In this context, we developed AptaMat, a simple and sensitive algorithm for ssNAs secondary structures comparison based on matrices representing the ssNAs secondary structures and a metric built upon the Manhattan distance in the plane. We applied AptaMat to several examples and compared the results to those obtained by the most frequently used metrics, namely the Hamming distance and the RNAdistance, and by a recently developed image-based approach. We showed that AptaMat is able to discriminate between similar sequences, outperforming all the other here considered metrics.

## Introduction

Single-stranded nucleic acids (ssNAs) are interesting molecules from both a biological and a biotechnological point of view. On one side, RNA is fundamental for protein synthesis and it has cellular structural, functional and regulatory roles. On the other side, both RNA and single -stranded DNA, in the form of aptamers, can be exploited as therapeutic or diagnostic tools or as biosensors [Kulabhusan *et al*., 2020]. Aptamers are, indeed, short single-stranded oligonucleotides able to bind a large variety of molecular targets with high specificity and dissociation constants in the nano- to picomolar range by adopting specific conformations [Li *et al*., 2020, Nimjee *et al*., 2017].

SsNAs function highly depends on their secondary (i.e. their base pairing pattern) and tertiary (i.e. their 3D organization) structures [Li *et al*., 2020, Mustoe *et al*., 2014, Nimjee *et al*., 2017], thus the computational prediction of these two levels of organization can help to understand ssNAs roles and interactions with other molecules. The prediction of the ssNAs secondary structures often precedes and guides the 3D modeling step and many tools have been developed at this scope ([Zuker., 2003, Gruber *et al*., 2008b, Sato *et al*., 2009]). The resulting output is usually a graphical representation of the predicted secondary structure (Figure 1c) and/or its dot-bracket notation (Figure 1b), which consists in a string of the same length as the sequence based on an alphabet of 3 characters: {“.”,”(“,”)”}. The symbol “.” indicates that the nucleotide in the corresponding position is unpaired, while “(“ and “)” correspond to the opening and closing positions of a base pair, respectively.

**Figure 1:**
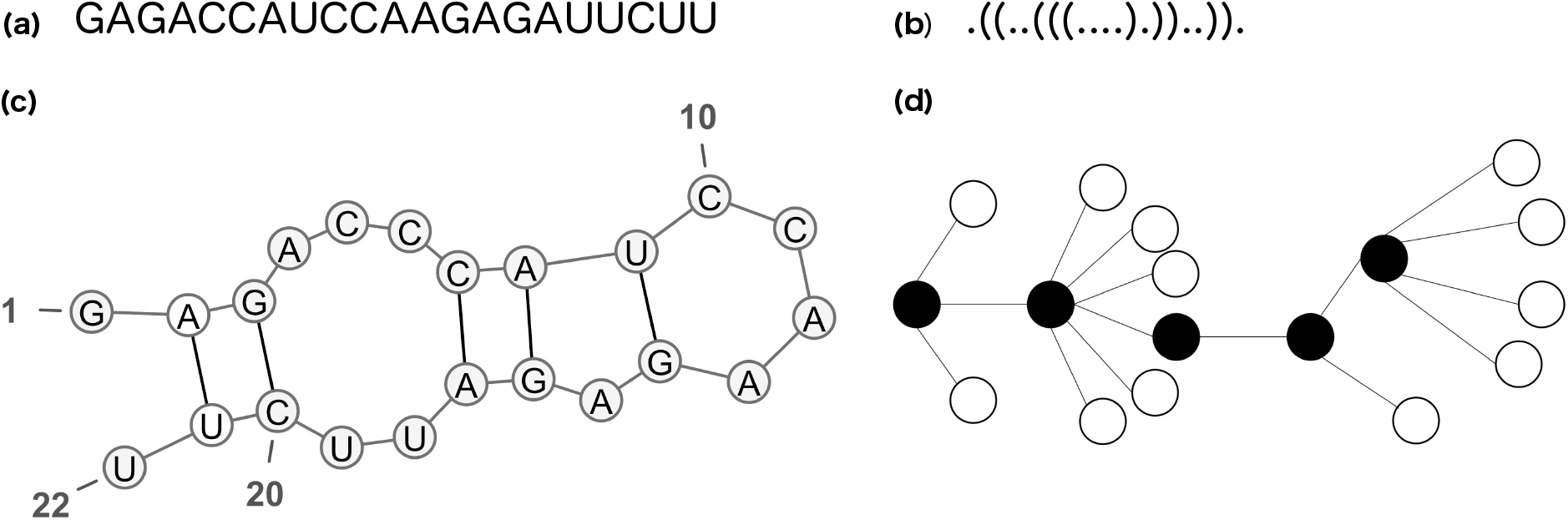
Example of representations of the secondary structure of sequence (**a**): dot-bracket notation (**b**), graphical representation realized with VARNA([Darty *et al*., 2009]) (**c**), and full tree representation (**d**).

The comparison of ssNAs secondary structures is a task as important as the prediction of the secondary structure itself. Comparing ssNAs structures can help to study the function and evolution of ssNAs, but also to design nucleotide sequences that fold into a given secondary structure and to predict mutations that can cause a conformational rearrangement. Therefore, different algorithms have been developed at this scope (see [Gruber *et al*., 2008a] for a review). Briefly, these can be classified in algorithms i) based on the minimum free energy [Washietl *et al*., 2005], ii) based on single structure [Shapiro *et al*., 1988, Moulton *et al*., 2000, Fontana *et al*., 1993, Flamm *et al*., 2001] and iii) considering the whole folding space [Hofacker *et al*., 1994, Bonhoeffer *et al*., 1993, Giegerich *et al*., 2004]. Among them, the most frequently applied are those working on single structures, such as the Hamming distance [Hamming., 1950] and the RNAdistance algorithm implemented in the ViennaRNA package [Hofacker *et al*., 2003]. The Hamming distance allows to compare two strings of the same length by counting the number of positions with different symbols. It is one of the simplest metrics used in the context of ssNAs, and it is usually calculated by counting the number of positions with different nucleotides (Equation 2). It can be adapted to strings in the dot-bracket notation, which is more suitable for secondary structures comparison. Conversely, RNAdistance is based on the comparison of ssNAs secondary structures represented as ordered rooted trees (Figure1d), deduced from the dot-bracket notation [Shapiro *et al*., 1988].

However, these two metrics sometimes fail in finding differences between secondary structures as showed in the example of Figure 2 adapted from [Ivry *et al*., 2009], where both the Hamming distance and RNAdistance cannot capture the differences between structures (b), (c) or (d) and the reference structure (a). Indeed, the Hamming distance only considers the total number of matching positions, without taking into account the correlations between the opening and closing positions, which are characteristic for the structure. On the other hand, RNAdistance works with a tree representation that, even at full resolution (i.e. without any loss of information with regard to the dot-bracket notation), might lead to an equivalent cost in the tree editing operations for structures that seem to have a different degree of proximity to the reference one. This is illustrated in Figure 2, and the details about the computation of RNAdistance can be found in Figure S1 of Supplementary Material. Interesting approaches for comparing ssNAs secondary structures based on image processing, such as DoPloCompare [Ivry *et al*., 2009], have been developed. These approaches consist in representing the secondary structures of the two compared ssNAs as dotplots and then processing them as images in order to measure the distance between the two structures. The use of dotplots allows to take into account the base pairs relative positions and it provides a finer description of the ssNA structure than RNAdistance [Ivry *et al*., 2009]. However, this approach can be laborious and sometimes it fails in finding the expected trend when comparing multiple structures to a reference one, as we will show later. Indeed, although the image processing approach is a novelty in the field, the proposed metrics use a combination of geometrical distance and histogram correlations that might hinder the nature of the proximity between the compared structures. Moreover, DoPloCompare seems to be not symmetric, which is an important requirement for many applications.

**Figure 2:**
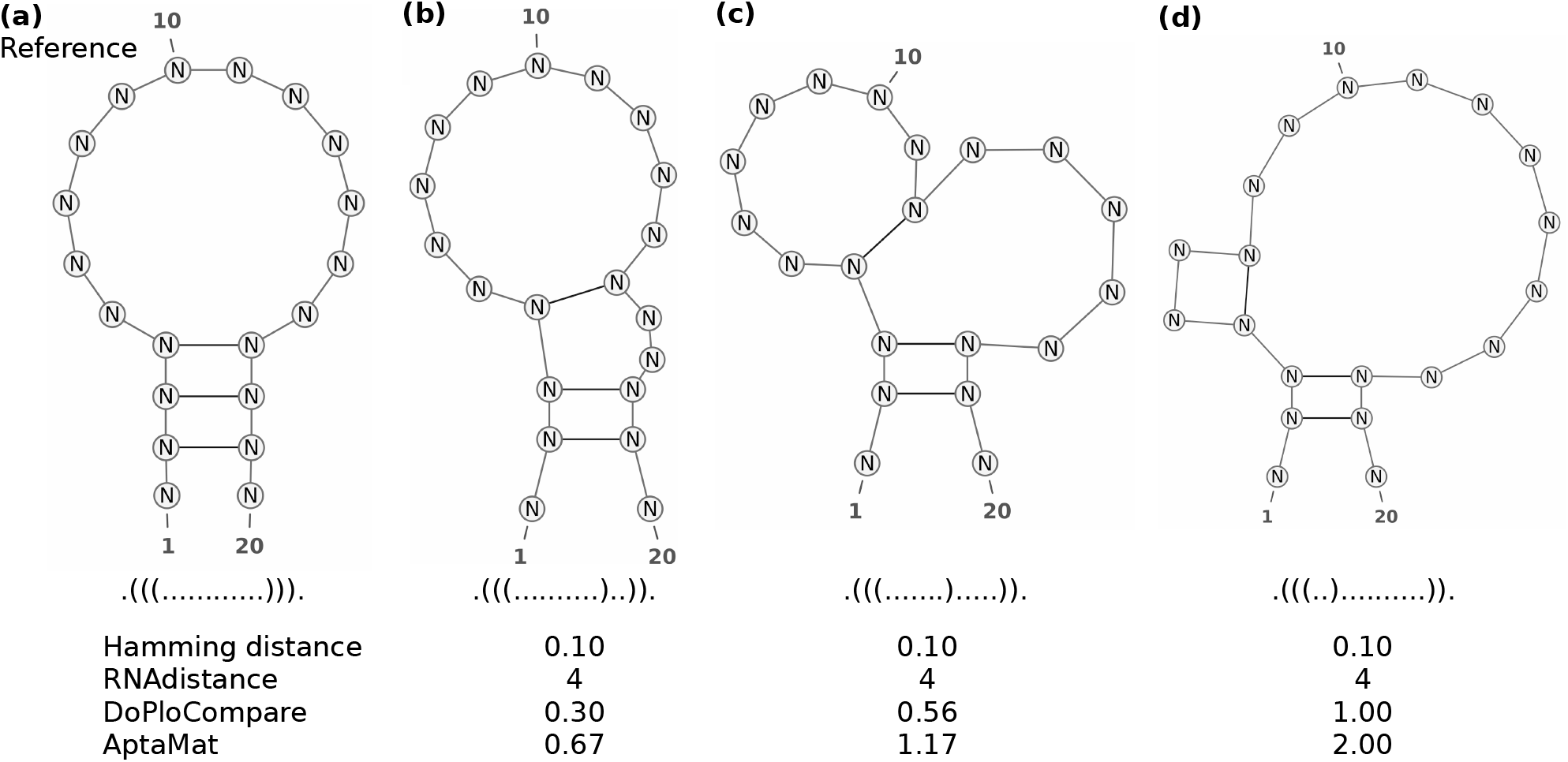
Reference (**a**) and alternative (**b, c**, and **d**) structures for ssNA 1. The Hamming, RNAdistance, DoPloCompare, and AptaMat distances are computed using structure (**a**) as reference.

Although there exist several other approaches to compare secondary structures, to our knowledge, none of them satisfy the desired properties: i) simple in terms of results interpretation; ii) easy to implement and to manipulate; iii) exploitable for the comparison of pairs of structures, but also of multiple structures to a reference one, and, most of all, iv) sensitive, in order to properly differentiate particularly close structures. Therefore, we developed a new algorithm, called AptaMat, which solves the issues of both the single structure-based and the image-based approaches. Briefly, AptaMat takes as input the secondary structure of two ssNAs (*S*_*A*_ and *S*_*B*_) of same length *L* in the dot-bracket notation and creates for each of them a matrix of size *L* × *L*, comparable to a dotplot with 1 and 0 instead of dots and blank cells, respectively. Indeed, the (*i, j*)^th^ entry of the matrix is either equal to 1 if the nucleotide in position *i* is paired with the nucleotide in position *j* or 0 if the nucleotides in positions *i* and *j* are not paired. For each base pair of each structure, we find the closest base pair on the other structure using the Manhattan distance between points in the plane. The distances between all the closest pairs are summed up and normalized by the total number of cells containing 1 in both matrices, in order to find the final AptaMat distance (Figures S2 and S3, Supplementary Material).

We applied our approach to i) 5 examples taken from the work by [Ivry *et al*., 2009] in order to make a direct comparison with the Hamming distance, RNAdistance and DoPloCompare and ii) to 5 structures of aptamers taken from the Protein Data Bank [Berman *et al*., 2000]. In addition, we *ad hoc* created an example capable of showing the advantages of our method as compared to both RNAdistance and the Hamming distance at the same time. The obtained results show that AptaMat is able to properly compare ssNAs secondary structures and to well discriminate among different structures. The python code implementing AptaMat is available on GitHub at https://github.com/GEC-git/AptaMat.git.

## Methods

### AptaMat algorithm

The AptaMat algorithm has been developed for the comparison and quantification of the differences between structures of pairs of ssNAs of the same length (*L*), with the main aim of investigating the effect of mutations on the ssNAs structure. The algorithm takes as input the two structures written in the dot-bracket notation, with one structure considered as reference. Starting from each input dot-bracket string a square matrix of *L* × *L* in size is created, where each matrix cell (*i, j*) corresponds to the position *i* of a nucleotide of the sequence relative to another position *j* of the same sequence. Therefore, each cell (*i, j*) contains either 1, if the nucleotide in position *i* is involved in a base pair with the nucleotide in position *j*, or 0 if not. The resulting matrices can be assimilated to dotplots, with 1 instead of a dot and 0 instead of blank cells. Although very simple, this representation allows to take into account the relative position of the base pairs in the ssNA sequence, thus retaining a more complete structural information as compared to the dot-bracket notation.

For the clarity of the algorithm description, we will call matrix *A* = (*a*_*ij*_) the one containing the information regarding the reference structure and matrix *B* = (*b*_*ij*_) the one storing the information of the structure we want to compare to the reference one. We want to define a distance between these matrices that reflects the proximity between cells containing 1 in both of them, i.e. those indicating a base pair. For this purpose, each matrix is embedded in the plane in the following way: each (*i, j*)^th^ entry that is equal to 1 is assimilated to the point with coordinates (*j, L* − *i* + 1). Hence, to a matrix representing a secondary structure we associate a set of points in the plane with coordinates in {1, …, *L*}^2^. Moreover, since both matrices are symmetrical, we consider only the entries below the diagonal. More precisely, let 𝒫_*A*_ := {(*j, L* − *i* + 1) ∈ ℕ ^2^ : *a*_*ij*_ = 1, 1 ≤ *j < i* ≤ *L*} be the set of points corresponding with structure *S*_*A*_. The set 𝒫_*B*_ is defined analogously. A natural way to measure the distance between the base pairs in the compared structures is to measure the distance between sets 𝒫_*A*_ and 𝒫_*B*_. At this scope, any distance between compact sets of points in ℝ^2^ could be appropriate for the method (e.g. Haussdorf distance [Huttenlocher *et al*., 1993]). At the moment, AptaMAT algorithm implements a metric based on the Manhattan distance, which was chosen for its simplicity, as it is expressed as the sum of the absolute differences between the coordinates of the compared points [Krause., 1988]. However, other distances can be easily implemented.

In AptaMat, for each point *P* in 𝒫_*A*_ we find the Manhattan distance to its nearest neighbor in 𝒫_*B*_, and vice versa. In order to handle all the differences between the structures, it is important to consider the distance in both directions (Figures S2 and S3, Supplementary Material). Indeed, both structures do not have necessarily the same number of base pairs. As a consequence, the distances in the two directions might not be the same and, more importantly, some base pairs might be excluded from the comparison. Therefore, considering only the distances in one direction might be source of mistake. Then, the shortest distances between 𝒫_*A*_ and 𝒫_*B*_ sets are summed up. Finally, the obtained distance is normalized by the total number of base pairs in structures *S*_*A*_ and *S*_*B*_. This is necessary because some distances might emerge twice in the calculation. Together with solving this issue, this sort of normalization gives a more important weight to base pairs in common between the two compared structures. The AptaMat distance, denoted by *D*_AM_ is, therefore, defined as

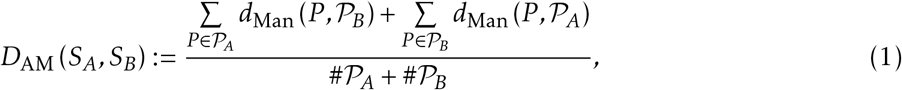

where, for any given point *P* = (*x, y*) ∈ ℝ^2^ and any finite subset 𝒞 ⊂ ℝ^2^, we denote by #𝒞 the cardinal of 𝒞, and by *d*_Man_(*P*, 𝒞) the Manhattan distance from *P* to its nearest neighbor in 𝒞.

We can easily check that *D*_AM_ is symmetric, and it is equal to 0 only when both structures are identical. In the light of this, the more the AptaMat distance is close to 0 the more the two compared structures are similar, independently on their length.

### Test set preparation

In order to confront AptaMat to the Hamming distance and RNAdistance in comparing ssNA secondary structures, we built a test set of 10 ssNA with known structures: 5 taken from the work by Ivry et al. [Ivry *et al*., 2009] and 5 taken from the PDB database (Table S1). The selected ssNA have different lengths (20 to 127 nucleotides) and different secondary structures, containing stems, hairpin/stem loops, bulges, internal loops and junctions. For each sequence, the reference secondary structure in the dot-bracket notation was either taken from [Ivry *et al*., 2009] or extrapolated using x3dna-dssr [Lu *et al*., 2003] and then used as the reference structure. In addition, for each sequence, 2 or more alternative structures where used to perform the comparison. The alternative structures for the examples taken from [Ivry *et al*., 2009] were obtained from the same article, while for those taken from the PDB database we used 6 different ssNA secondary structure prediction tools, namely Mfold [Zuker., 2003], LinearFold [Huang *et al*., 2019], CentroidFold [Hamada *et al*., 2009], RNAfold [Gruber *et al*., 2008a], RNAstructure [Reuter *et al*., 2010] and MC-Fold [Parisien *et al*., 2008] to obtain at least two different secondary structures for each ssNA. This was achieved when the prediction tools were not able to correctly predict the secondary structure of the processed sequences. In addition, we *ad hoc* designed an additional example to clearly show the advantages of AptaMAT over the two selected metrics of comparison. At this scope, we designed critical secondary structures able to highlight the limits of the other metrics and the strengths of AptaMat.

### Comparison methods

We compared AptaMat to two of the most used methods of ssNAs secondary structures comparison: the Hamming distance ([Hamming., 1950]) and RNAdistance from the ViennaRNA package [Hofacker *et al*., 2003]. The former computes the distance between two ssNAs structures of same length *L*, by calculating

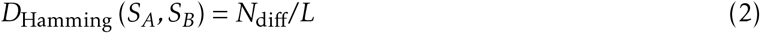

where *N*_diff_ is the number of unmatched positions between the two strings corresponding to the the dot-bracket notation of the compared structures. RNAdistance computes the distance between two ssNAs structures by representing them as ordered rooted trees. At a full resolution, this representation is deducible from the dot-bracket notation by assigning each unpaired nucleotide to a leaf and each base pair to an internal node, as showed in Figure 1d. In order to calculate the distance between two trees, the tree editing approach is used, which consists in a series of edit operations (deletion, insertion or mutation of a node), to which a cost is assigned and that allow to transform a tree *T*_*A*_ into a tree *T*_*B*_. The resulting distance *D*_RNA_(*S*_*A*_, *S*_*B*_) corresponds to the minimal total cost of the series of operations allowing to transform one tree into the other.

In addition, for the structures taken from [Ivry *et al*., 2009] (Table S1), we included in the benchmark of AptaMat the comparison with the algorithm DoPloCompare, which uses an approach based on image processing to measure the distance between two ssNAs secondary structures. This algorithm has been selected for comparison with AptaMat, because of its higher sensitivity as compared to the Hamming distance and RNAdistance (Figure 2), and because it is based on the dotplot diagrams of the compared structures, as AptaMat. The distance grade proposed in this algorithm to compare two structures *S*_*A*_ and *S*_*B*_ can be defined as

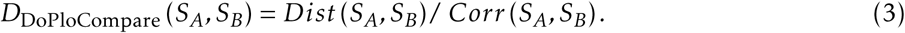

The *Dist* (*S*_*A*_, *S*_*B*_) term corresponds to the geometrical distance from the points in the dotplot diagram of structure *S*_*A*_ (reference) to the dotplot diagram of structure *S*_*B*_ (alternative). The *Corr* term is related to the cross correlation between histogram vectors built from the dotplot diagrams of both structures by adding the number of points in four different directions (X, Y, diagonal and antidiagonal). Although the Dist term in DoPloCompare is somehow similar to AptaMat, it doesn’t seem to be symmetrically defined, and hence it does not take into account the number of base pairs in the alternative structure. On the other hand, the *Corr* term accounts for the similarity in the order and number of elements that both structures contain, even if the base pairs involved in these elements are not the same in structures *S*_*A*_ and *S*_*B*_.

## Results and Discussion

We used AptaMat to measure the distance between pairs of secondary structures using the ssNAs reported in Table S1 and we compared the AptaMat distance with the Hamming distance and RNAdistance. Among these, for ssNAs 2, 4, 5, and 7 (Figures 5, 3 and Figures S5 and S6) the Hamming distances, RNAdistances and AptaMat distances of the alternative secondary structures from the reference one follow the same trend. This shows the coherence between our method and the most used distance metrics when there is a clear difference between the compared secondary structures in terms of both dot-bracket notation and the trees used to calculate RNAdistance. We discuss here the results for ssNA 7 (Table S1 and Figure 3), since for this ssNA we could gather 3 different alternative structures, which allows for a more extensive analysis. The three distances from the reference structure **(a)** progressively increase proceeding from the alternative structure **(b)**, obtained by RNAstructure [Reuter *et al*., 2010] (Hamming distance = 0.15, RNAdistance = 8 and AptaMat distance = 0.44), to **(d)**, obtained by RNAfold ([Gruber *et al*., 2008a]) (Hamming distance = 0.40, RNAdistance = 32 and AptaMat distance = 9.95). Indeed, the reference secondary structure (a) made of a stem, a multibranched loop, a bulge and two hairpin/stem loops is progressively lost. The alternative structure **(a)** is close to the reference: instead of the original G9-C20 base pair, it has a base pair between C7 and G17 and one between A8 and T18. This leads to the transformation of the bulge in an internal loop and the reduction of the width of the multi-branched loop. Structure **(c)** has a much wider multi-branched loop because of the loss of 5 base pairs, which also shorten the two hairpin/stem loops, with one of them becoming a bulge. Finally, structure **(d)** only conserves 2 hairpin/stem loops and the bulge but they do not involve the same positions as in the reference.

**Figure 3:**
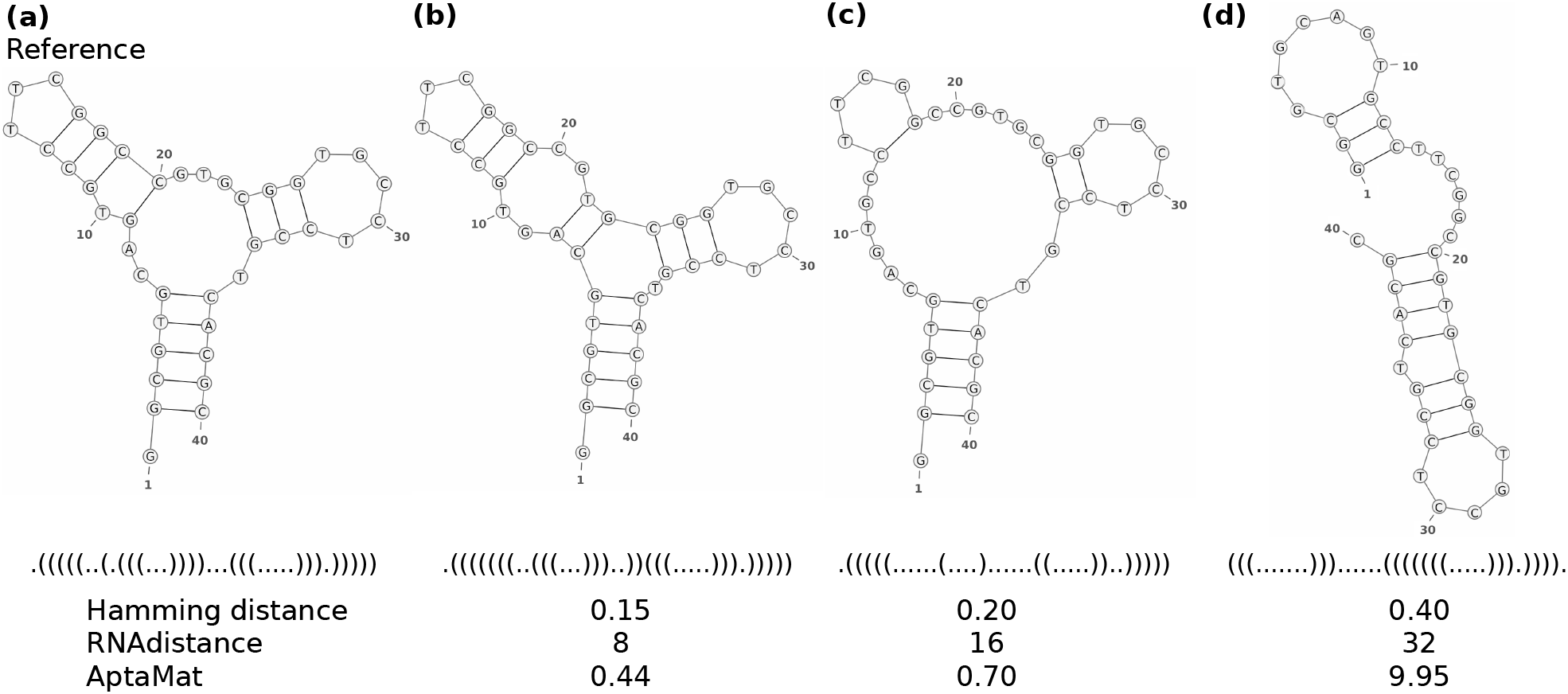
SsNA 7 shows the ability of AptaMat in comparing ssNAs secondary structures. The three metrics (Hamming distance, RNAdistance and AptaMat) indicate that the alternative structures (**b**), (**c**) and (**d**) are progressively farther from the reference secondary structure (**a**).

However, sometimes the structural differences between two ssNAs are quite subtle and the Hamming distance and RNAdistance are not able to discriminate between structures. A striking example is represented by ssNA 1 (Table S1 and Figure 2), which has been taken from [Ivry *et al*., 2009]. This example is not based on the analysis of a proper ssNA sequence but it focuses directly on structures. As shown in Figure 2, the three structures compared to the reference differ from this latter and one from another. The three alternative structures have an additional bulge, which becomes progressively wider from structure **(b)** to structure **(d)**, since the third base pair progressively shifts towards the 5’ end. However, both the Hamming distance and RNAdistance predict the same distance to the reference for the three alternative structures. Indeed, the Hamming distance counts the number of mismatches between the dot-bracket strings to compare. Therefore, it doesn’t take into account the position of the nucleotides involved in base pairs. As a result, any information about the structure is lost and different secondary structures with the same number of mismatching positions as compared to a reference structure will have the same Hamming distance from it. In ssNA 1 all the alternative structures have 2 mismatching positions, which, accordingly to Equation 2, leads to a Hamming distance of 0.10 in all the cases. Conversely, RNAdistance takes into account the correlation between opening and closing position of the dot-bracket notation strings. However, it might happen that the series of editing operations of two comparisons have an equivalent weight leading to the same RNAdistance, as it occurs in the example of Figure 2 (see Figure S1 for the details). On the opposite, both AptaMat and DoPloCompare are able to correctly calculate the distance trend, with the first alternative structure being the closest to the reference (AptaMat distance = 0.67 and DoPloCompare distance = 0.30) and the third alternative structure being the furthest (AptaMat distance = 2.00 and DoPloCompare distance = 1.00).

SsNAs 3, 6, 9, and 10 also show the same RNAdistance and/or Hamming distance between different predicted structures and their reference (Figures S4, S7, S9 and S10). As mentioned before, the Hamming distance will be the same if the alternative structures have the same number of mismatching positions as compared to the reference one. However, depending on the number and the position of the mismatches, the structural difference might become highly relevant and lead to wrong conclusions about the similarity of a structure to a reference one. In order to highlight the issues arising from the Hamming distance and RNAdistance in a unique example, we *ad hoc* created the example reported in Figure 4 (ssNA 11 in Table S1). As for ssNA 1, we decided to focus on the secondary structures and not on the nucleotide sequence. The structures (b) and (c) have the same Hamming distance to the reference structure (a), since they both have 4 mismatching positions. However, structure (c) doesn’t have the N12-N19 and N13-N18 base pairs, leading to the loss of the hairpin/stem loop. Conversely, structure (b) maintains the reference structure consisting of a hairpin, a bulge, an internal loop and the hairpin/stem loop, although the bulge is 3 nucleotides shorter and the internal loop 3 nucleotides wider. This clearly comes out from RNAdistance and AptaMat, both indicating that structure **(b)** is closer to the reference structure than structure **(c)**.

**Figure 4:**
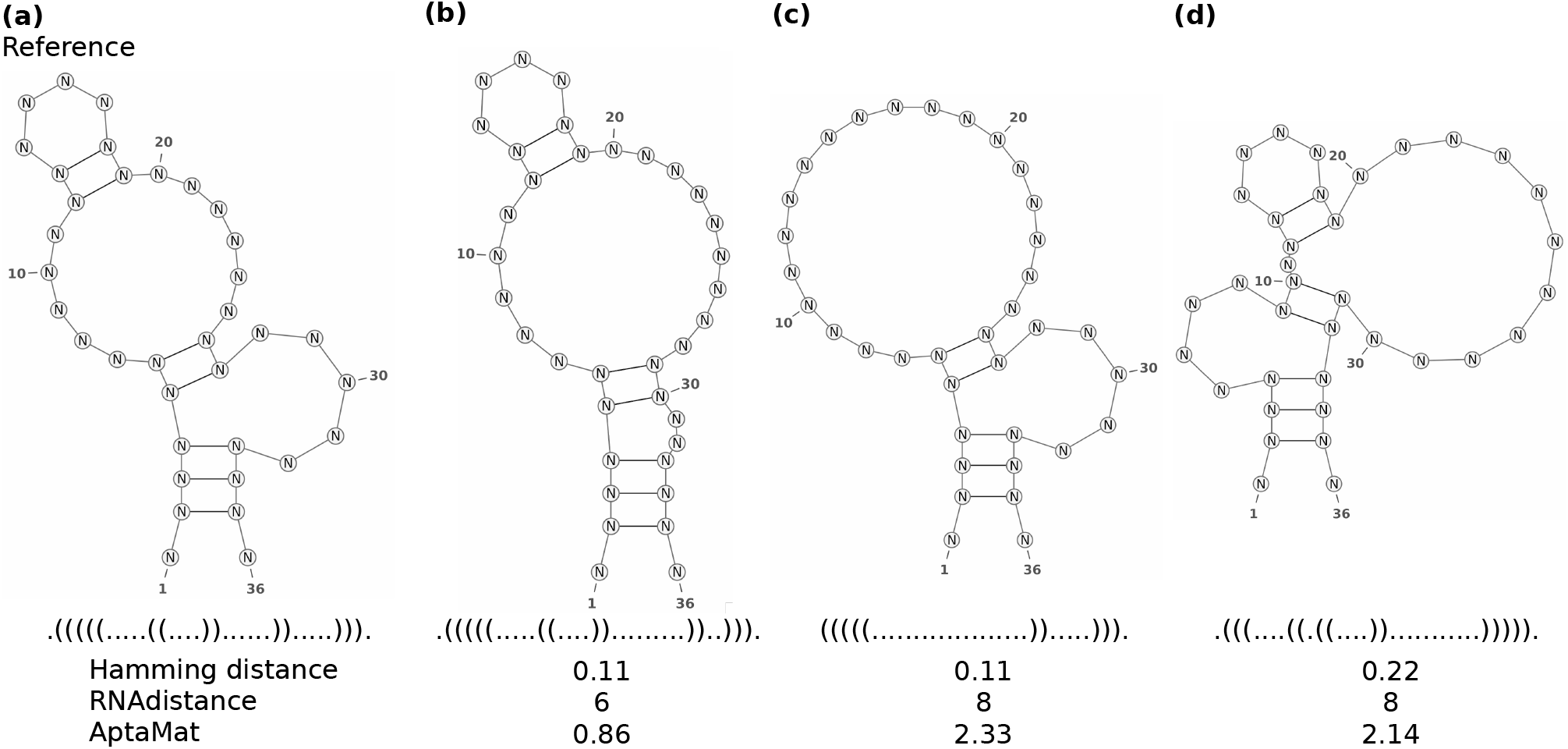
SsNA 11 shows the limits of the Hamming distance and RNAdistance in comparing ssNAs secondary structures. Alternative structures (**b**) and (**c**) have the same Hamming distance to the reference secondary structure (**a**), although structure (**c**) misses the hairpin/stem loop. Alternative structures (**c**) and (**d)** have the same RNAdistance to the reference secondary structure (**a**), although the bulge and the internal loop involve different nucleotides as compared to the reference.

Within the example in Figure 4 we can further investigate the limits of RNAdistance, since structures **(c)** and **(d)** have the same RNAdistance to the reference structure **(a)**. Indeed, the sum of the weights associated to the editing tree operations from **(c)** to **(a)** and from **(d)** to **(a)** is the same (Figure S10). Conversely, although both the alternative structures **(c)** and **(d)** are far from the reference, AptaMat indicates that structure **(d)** is slightly closer to the reference than structure **(c)**. As previously said, because of the loss of the missing N12-N19 and N13-N18 base pairs, structure **(c)** doesn’t have the hairpin/stem loop present in the reference structure, although the hairpin and the bulge involve the same nucleotides as in the reference (N2-N35, N3-N34, N4-N33, N5-N27 and N6-N26). Conversely, structure **(d)** keeps the overall structure of the reference and the same number of base pairs, but the bulge and the internal loop don’t involve exactly the same nucleotides as the reference: base pairs N2-N35, N3-N34, N4-N33, N12-N19 and N13-N18 are maintained, while base pairs N5-N27 and N6-N26 are replaced by base pairs N9-N22 and N10-N21. Together with being able to observe even a slight difference in the distance from structures **(c)** and **(d)** to the reference structure **(a)**, AptaMat focuses more on the overall secondary structure and the conserved base pairs than on the matching positions of the dot-bracket notations, as required when working on ssNAs, whose function is structure-dependent. Similar observations can be done for ssNAs 9 and 10 (Figures S9 and S10), where the ssNA reference secondary structure has been extrapolated from the 2VJU and 5HRU PDB entries, respectively.

Together with being able to distinguish between differences in pairs of compared structures, AptaMat is capable to establish more meaningful ranking of the alternative secondary structures in terms of distance from the reference as compared to the Hamming distance and RNAdistance in all the examples herein presented. This is important when investigating the effect of sequence mutations on the ssNAs secondary structure. In this context, ssNAs 3, 5, 6, 8 and 9 (Table S1) show the limits of these latter methods as compared to AptaMat. Here we focus our discussion on ssNA 6, which has more alternative structures than ssNAs 3, 5 and 9, and more subtle modifications than ssNA 8. Thus, this example offers the possibility to deeply explore the differences between the considered metrics. SsNA 6 (PDB ID: 1NGO) has a simple hairpin/stem loop structure (Figure S7). The alternative structure **(b)** obtained by CentroidFold is correctly considered by the three metrics as the closest one to the experimental structure (Hamming distance = 0.074, RNAdistance = 2 and AptaMat = 0.091). AptaMat then indicates that the alternative structure **(d)** obtained by MC-Fold is closer to the reference (AptaMat distance = 0.20) than the alternative structure **(c)** obtained by RNAfold (AptaMat distance = 0.22), since the former only misses two pairs of bases (T5-G23 and T6-G22) while maintaining the overall structure. Conversely, structure (c) has 2 additional base pairs that lead to the loss of the characteristic loop of 1NGO (Figure S7). On the opposite, the Hamming distance fails in finding this difference, and RNAdistance suggests the opposite trend, with structures (c) and (d) having an RNAdistance of 6 and 8, respectively. Similar conclusions are applicable to ssNA 3 and 8 (Figures S4 and S8), while for ssNAs 5 and 9 (Figures S6 and S9) the Hamming distance indicates an opposite and inadequate ranking of the two alternative structures in terms of distance from the reference, because of the different number of mismatches.

The overall better performance of AptaMat as compared to the Hamming distance and RNAdistance in ranking the alternative secondary structures in terms of distance from a reference is particularly evident for structures having a similar distance from the reference, which are more difficult to properly rank. The ability of AptaMat in doing so is due to the higher weight given by our algorithm to the relative position of the base pairs. This leads to focus on the global secondary structure more than on the local differences from the reference secondary structure. As previously mentioned, this is of particular importance for the comparison of ssNAs, since their function highly depends on their global 3D structure and only to a minor extent on local sequence information.

In addition, together with the better performance as compared to RNAdistance and the Hamming distance, AptaMat has the advantage of being easy to interpret. Indeed, by observing the herein reported examples, we could suggest a threshold of about 2 to conclude on the proximity of a sequence to the reference one: an AptaMat distance below this threshold indicates that the two structures are close, while a greater distance indicates that the two structures are far one from another. This is supported also by a benchmark study on the the available ssNAs secondary structures prediction tools we performed (article in preparation), but this threshold can be adapted for different applications. On the opposite, RNAdistance relies on tree editing operations with fixed weights, which cannot be interpreted in an absolute way: although the lower is the RNAdistance the closer are the compared structures, an RNAdistance of 8 might indicate close structures as in ssNA 7 (Figure 3b) but it can also be associated to more relevant changes in the ssNA structures as in ssNA 11 (Figure 4c).

The analysis of the alternative structures ranking relative to the reference structure allows also to highlight the limits of DoPloCompare as compared to AptaMat. SsNAs 2, 4 and 5 (Figure 5 Figures S5 and S6) have a DoPloCompare trend opposite not only to AptaMat but also to the Hamming distance and RNAdistance. We argue that this is due to the *Corr* term in DoPloCompare, which, as we mentioned before, accounts for the similarities in the number and order of the elements (stems, loops, etc.) in the compared structures. In the three previous examples, the structures that are found to be closer to the reference one are those having a more similar number of elements, despite the fact that the base pairs involved in these elements are not the same. For example, if we consider ssNA 2 (Figure 5), we can clearly see that the alternative structures **(b)** and **(c)** are both structurally far from the reference structure **(a)**. However, the structure **(b)** is closer to the reference **(a)** (Hamming distance = 0.15, RNAdistance = 24 and AptaMat = 6.35) than the alternative structure **(c)** (Hamming distance = 0.41, RNAdistance = 26 and AptaMat = 7.50), as correctly indicated by the Hamming distance, RNAdistance and AptaMat. Indeed, structure **(b)** maintains the secondary structure of the reference except for 3 missing base pairs (G28-C37, G29-C36 and C30-G35), while structure **(c)** has 4 additional base pairs (C5-G39, C6-G38, C12-G27, U13-G26), leading to a significant change in the global structure. DoPloCompare indicates that this latter structure is closer to the reference (DoPloCompare = 0.12) than structure (b) (DoPloCompare = 0.13), because structure **(c)** has two hairpin/stem loops and an internal loop as structure **(a)**, while structure **(b)** only has a hairpin/stem loop and and an internal loop. However, the global structure **(c)** differs from those in structure **(a)**, because of a different base pairs pattern. In addition, the DoPloCompare scores are close to 0, suggesting a high similarity of the alternative structures to the reference one, which is clearly not the case as indicated by RNAdistance and AptaMat. Similar observations can be done for ssNAs 4 and 5 (Figures S5 and S6). Furthermore, looking at the DoPloCompare scores obtained for ssNAs 1 to 5, it seems that they depend on the sequence length: although the alternative structures of ssNAs 1 (Figure 2) are globally close to the reference one, they show a DoPloCompare score which is higher than those obtained for ssNAs 2 to 5, where the alternative structures are very far from the reference, as also showed by the RNAdistance and AptaMat.

**Figure 5:**
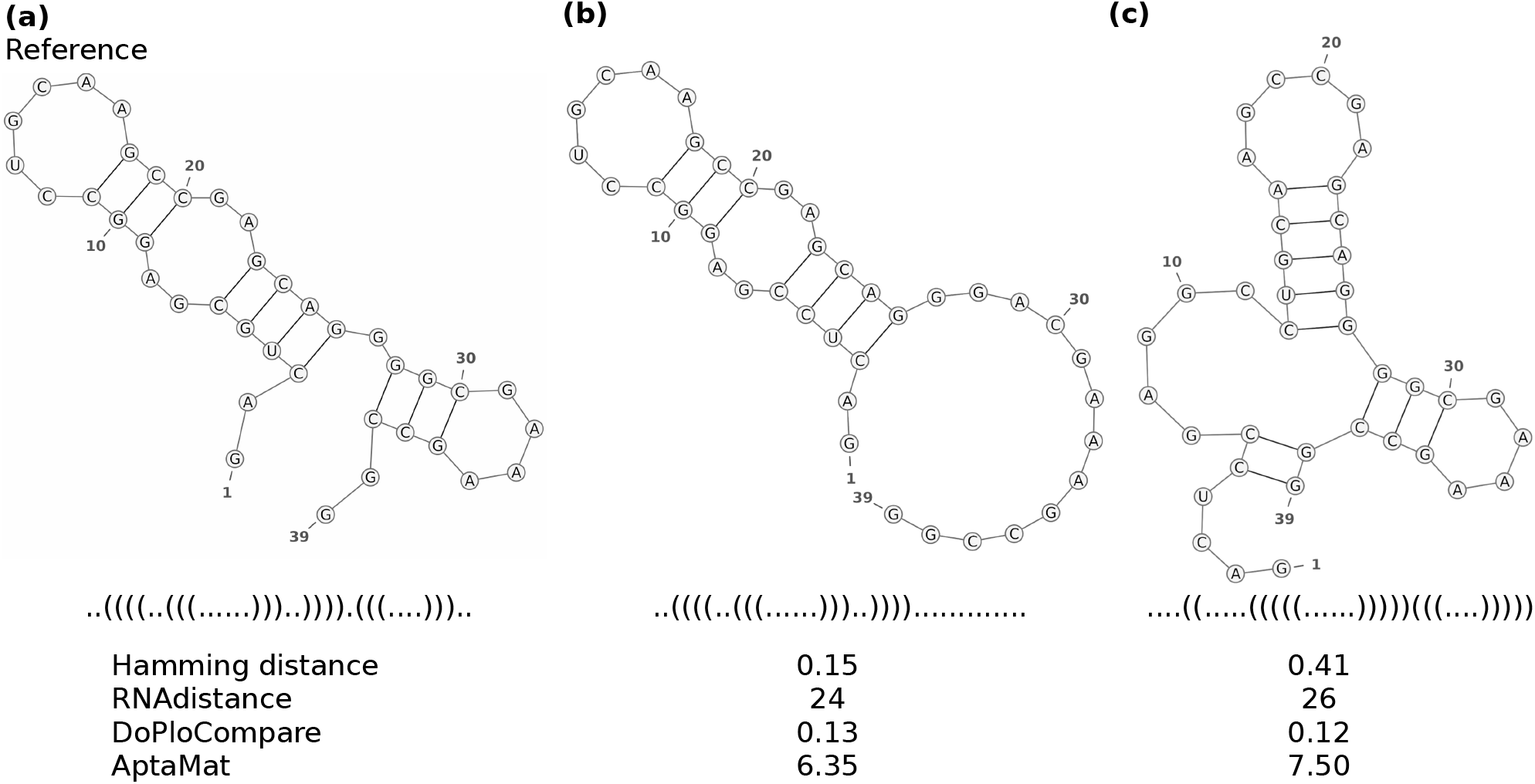
SsNA 2 shows the limits of DoPloCompare in ranking the alternative secondary structures in terms of distance from the reference. The alternative structure (**b**) is closer to the reference according to the Hamming distance, RNAdistance and AptaMat, since it has the same internal loop and one hairpin/stem loop, while the alternative structure (**c**) involves different nucleotides in one of the hairpin/stem loops and it assumes a 3-ways junction structure.

## Conclusion

Being able to compare ssNAs secondary structures is fundamental to understand the function and evolution of this kind of biomolecules, to design ssNAs with a desired secondary structure or even to predict the conformational effects of sequence mutations. In the light of this, in this work we present AptaMat, a new matrix-based algorithm, capable of comparing pairs of ssNAs secondary structures of the same length *L*. AptaMat takes as input the two ssNAs structures in the dot-bracket notation and, for each of them, creates a matrix of size *L* × *L*, named *A* = (*a*_*ij*_) and *B* = (*b*_*ij*_). The (*i, j*)^th^ entry of the matrix is either equal to 1 if the nucleotide in position *i* is paired with the nucleotide in position *j* or 0 if the nucleotides in positions *i* and *j* are not paired. Then, for each 1 ≤ *i < j* ≤ *L* such that *a*_*ij*_ = 1, the Manhattan distance to the closest entry equal to 1 in matrix *B*, and vice versa, is calculated. The distances between all the closest pairs are summed up and normalized by the total number of cells containing 1 in both matrices, leading to AptaMat distance.

We compared AptaMat to two of the most used metrics for ssNAs secondary structures comparison, namely the Hamming distance and RNAdistance, and to a more recent approach based on image processing, DoPloCompare, by [Ivry *et al*., 2009]. In order to do this, we chose 5 structures taken from the examples reported in the work by Ivry et al. and 5 structures taken from the PDB database. In addition, we *ad hoc* created an additional structure in order to clearly show the advantages of AptaMat over the Hamming distance and RNAdistance.

We showed that AptaMat is able to properly distinguish between different structures, presenting a higher sensitivity as compared to the Hamming distance and RNAdistance. In addition, our method allows to more adequately rank the ssNAs structures as a function of their distance from a reference in all the examples herein discussed, which is not the case for the Hamming distance, RNAdistance and DoPloCompare. Moreover, it is easy to interpret, with an AptaMat distance of 2 as a reasonable threshold between close and far structures, but this threshold can be adapted depending on the applications. By definition, AptaMat is less affected by ssNA length than other of the considered metrics. Additionally, AptaMat is easy to implement and to manipulate. Indeed, we plan to extend its usage to ssNAs of different lengths by previous alignment, and to peculiar structures, such as pseudoknots and G-quadruplex, which represent a challenging task in nucleic acids modeling.

## Data Availability

The python code for AptaMat is available at https://github.com/GEC-git/AptaMat.git

## References

Berman, Helen M. and Westbrook, John and Feng, Zukang and Gilliland, Gary and Bhat, T. N. and Weissig, Helge and Shindyalov, Ilya N. and Bourne, Philip E. (2000). The Protein Data Bank. Nucleic Acids Res, 28(3), 235–242.

Bonhoeffer, S. and McCaskill, J. S. and Stadler, P. F. and Schuster, P. (1993). RNA multi-structure landscapes - A study based on temperature dependent partition functions. European Biophysics Journal, 22(1), 13–24.

Darty, K., Denise, A., and Ponty, Y. (2009). VARNA: Interactive drawing and editing of the RNA secondary structure. Bioinformatics, 25(15), 1974–1975.

Flamm, C., Hofacker, I. L., Maurer-Stroh, S., Stadler, P. F., and Zehl, M. (2001). Design of multistable RNA molecules. RNA, 7(2), 254–265.

Fontana, W., Konings, D. A., Stadler, P. F., and Schuster, P. (1993). Statistics of RNA secondary structures. Biopolymers, 33(9), 1389–1404.

Giegerich, R., Voß, B., and Rehmsmeier, M. (2004). Abstract shapes of RNA. Nucleic Acids Research, 32(16), 4843–4851.

Gruber, A. R., Bernhart, S. H., Hofacker, I. L., and Washietl, S. (2008). Strategies for measuring evolutionary conservation of RNA secondary structures. BMC bioinformatics, 9(1), 122.

Gruber, A. R., Lorenz, R., Bernhart, S. H., Neuböck, R., and Hofacker, I. L. (2008). The Vienna RNA websuite. Nucleic acids research, 36(Suppl2), W70–W74.

Hamada, M., Kiryu, H., Sato, K., Mituyama, T., and Asai, K. (2009). Prediction of RNA secondary structure using generalized centroid estimators. Bioinformatics,25(4), 465–473.

Hamming, R. W. (1950). Error Detecting and Error Correcting Codes. Bell System Technical Journal, 29(2), 147–160.

Hofacker, I. L. (2003). Vienna RNA secondary structure server. Nucleic Acids Research, 31(13), 3429–3431.

Hofacker, I. L., Fontana, W., Stadler, P. F., Bonhoeffer, L. S., Tacker, M., and Schuster, P. (1994). Fast folding and comparison of RNA secondary structures. Monatshefte für Chemie Chemical Monthly, 125(2), 167–188.

Huang, L., Zhang, H., Deng, D., Zhao, K., Liu, K., Hendrix, D. A., and Mathews, D. H. (2019). LinearFold: Linear-time approximate RNA folding by 5’-to-3’ dynamic programming and beam search. Bioinformatics, 35(14), i295–i304.

Huttenlocher, D. P., Klanderman, G. A., and Rucklidge, W. J. (1993). Comparing Images Using the Hausdorff Distance. IEEE Transactions on Pattern Analysis and Machine Intelligence, 15(9), 850–863.

Ivry, T., and Michal, S., Avihoo, A., Sapiro, G., Barash, D. (2009). An image processing approach to computing distances between RNA secondary structures dot plots. Algorithms for Molecular Biology, 4, 4.

Krause, E. (1988). Taxicab Geometry: An Adventure in Non-Euclidean Geometry, 72. Dover Publications.

Kulabhusan, Prabir Kumar and Hussain, Babar and Yüce, Meral (2020). Current perspectives on aptamers as diagnostic tools and therapeutic agents. Pharmaceutics, 12(7), 1–23.

Li, Long and Xu, Shujuan and Yan, He and Li, Xiaowei and Yazd, Hoda Safari and Li, Xiang and Huang, Tong and Cui, Cheng and Jiang, Jianhui and Tan, Weihong (2020). Nucleic Acid Aptamers for Molecular Diagnostics and Therapeutics: Advances and Perspectives. Angewandte Chemie International Edition, 59(5), 2–13.

Lu, X. and Olson, W. K. (2003). 3DNA: a software package for the analysis, rebuilding and visualization of three-dimensional nucleic acid structures. Nucleic Acids Research, 31(17), 5108–5121.

Moulton, V., Zuker, M., Steel, M., Pointon, R., and Penny, D. (2000). Metrics on RNA Secondary Structures. Journal of Computational Biology, 7(1-2), 277–292.

Mustoe, A. M., Brooks, C. L., and Al-Hashimi, H. M. (2014). Hierarchy of RNA Functional Dynamics. Annual Review of Biochemistry, 83(1), 441–466.

Nimjee, Shahid M. and White, Rebekah R. and Becker, Richard C. and Sullenger, Bruce A. (2017). Aptamers as Therapeutics. Annual Review of Pharmacology and Toxicology, 57(1), 61–79.

Parisien, M. and Major, F. (2008). The MC-Fold and MC-Sym pipeline infers RNA structure from sequence data. Nature, 452(7183), 51–55.

Reuter, J. S. and Mathews, D. H. (2010). RNAstructure: Software for RNA secondary structure prediction and analysis. BMC Bioinformatics, 11(1), 1–9.

Sato, K., Hamada, M., Asai, K., and Mituyama, T. (2009). CentroidFold: A web server for RNA secondary structure prediction. Nucleic Acids Research, 37(SUPPL. 2), W277–W280.

Shapiro, B. A. (1988). An algorithm for comparing multiple RNA secondary structures. Bioinformatics, 4(3), 387–393.

Washietl, S., Hofacker, I. L., and Stadler, P. F. (2005). Fast and reliable prediction of noncoding RNAs. Proceedings of the National Academy of Sciences of the United States of America, 102(7), 2454–2459.

Zuker, M. (2003). Mfold web server for nucleic acid folding and hybridization prediction. Nucleic Acids Research, 31(13), 3406–3415.

