## Supplemental Material for "AptaMat: a matrix-based algorithm to compare single-stranded oligonucleotides secondary structures"

### Supplementary Material for 'AptaMat: a matrix-based algorithm to compare single stranded oligonucleotides secondary structures'

Thomas Binet[1], Bérangère Avalle[1], Miraine Dávila Felipe[2,*] and Irene Maffucci[1,*]
[1]Université de technologie de Compiègne, UPJV, CNRS, Enzyme and Cell Engineering, Centre de recherche Royallieu - CS 60 319 - 60 203 Compiègne Cedex
[2]Université de technologie de Compiègne, LMAC (Laboratory of Applied Mathematics of Compiègne), CS 60 319 - 60 203 Compiègne Cedex
*To whom correspondence should be addressed.

#### Contents

#### List of Tables

Table S1 ssNAs test set nomenclature . . . . . . . . . . S2

#### List of Figures

Figure S1 Tree Representation . . . . . . . . . . S2
Figure S2 AptaMat flowchart . . . . . . . . . . S3
Figure S3 AptaMat algorithm . . . . . . . . . . S4
Figure S4 Results for ssNA 3 . . . . . . . . . . S5
Figure S5 Results for ssNA 4 . . . . . . . . . . S5
Figure S6 Results for ssNA 5 . . . . . . . . . . S6
Figure S7 Results for ssNA 6 . . . . . . . . . . S7
Figure S8 Results for ssNA 8 . . . . . . . . . . S7
Figure S9 Results for ssNA 9 . . . . . . . . . . S8
Figure S10 Results for ssNA 10 . . . . . . . . . . S8
Figure S11 Details of ssNA 2 . . . . . . . . . . S9

Table S1: ssNAs test set nomenclature.

| No. | Code PDB[a] | Length | Reference dot-bracket notation |
|---|---|---|---|
| ssNAs from [Ivry *et al.*, 2009] | | | |
| 1 | NA | 20 | .(((............))). |
| 2 | NA | 39 | ..((((..(((......)))..)))).(((....))).. |
| 3 | NA | 43 | .((((((((((((...((.......))...))))))))).))) |
| 4 | NA | 56 | ....((((.(((((.......))))).))))....((((........)))).)))) |
| 5 | NA | 127 | (((((((...((((((((((((((..((((((.(((((((.(((.((....((((..((((... ...))))..))))))))))))))))))).)).)))).)))))).))))))))).)))))))) |
| ssNAs from PDB database | | | |
| 6 | 1NGO | 26 | (((((((((((.....)))))))))) |
| 7 | 3HXO | 40 | .(((((..(.(((...))))...(((.....))).))))) |
| 8 | 1SNJ | 36 | (((((((((((..))))((((..))))..))))))) |
| 9 | 2VJU | 35 | ((..((((((((....)))).))))......)).. |
| 10 | 5HRU | 32 | ((((.........(((((.....))))))))) |
| *Ad hoc* designed ssNA | | | |
| 11 | NA | 36 | .(((((.....((....))......)).....))). |

[a]The PDB code is not available for the structures taken from [Ivry *et al.*, 2009] and for the *ad hoc* designed one.

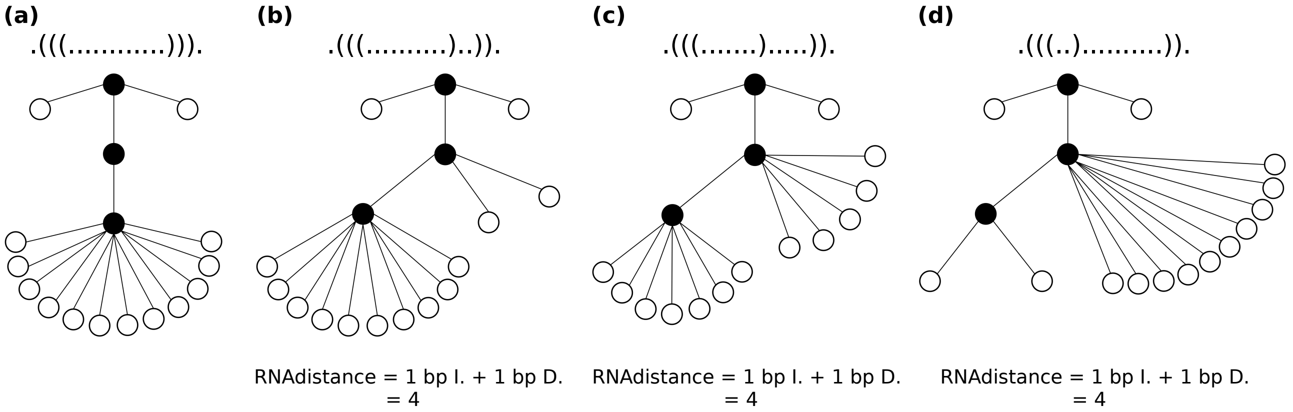


Figure S1: Dot-bracket and tree representations of ssNA 1 **(a)**, and the alternative structures **(b)**, **(c)**, and **(d)**. The computation of RNAdistance for each of the alternative structures is provided below the corresponding trees. Here “bp” means base pair, “I.” corresponds to an insertion operation, and “D.” to a deletion

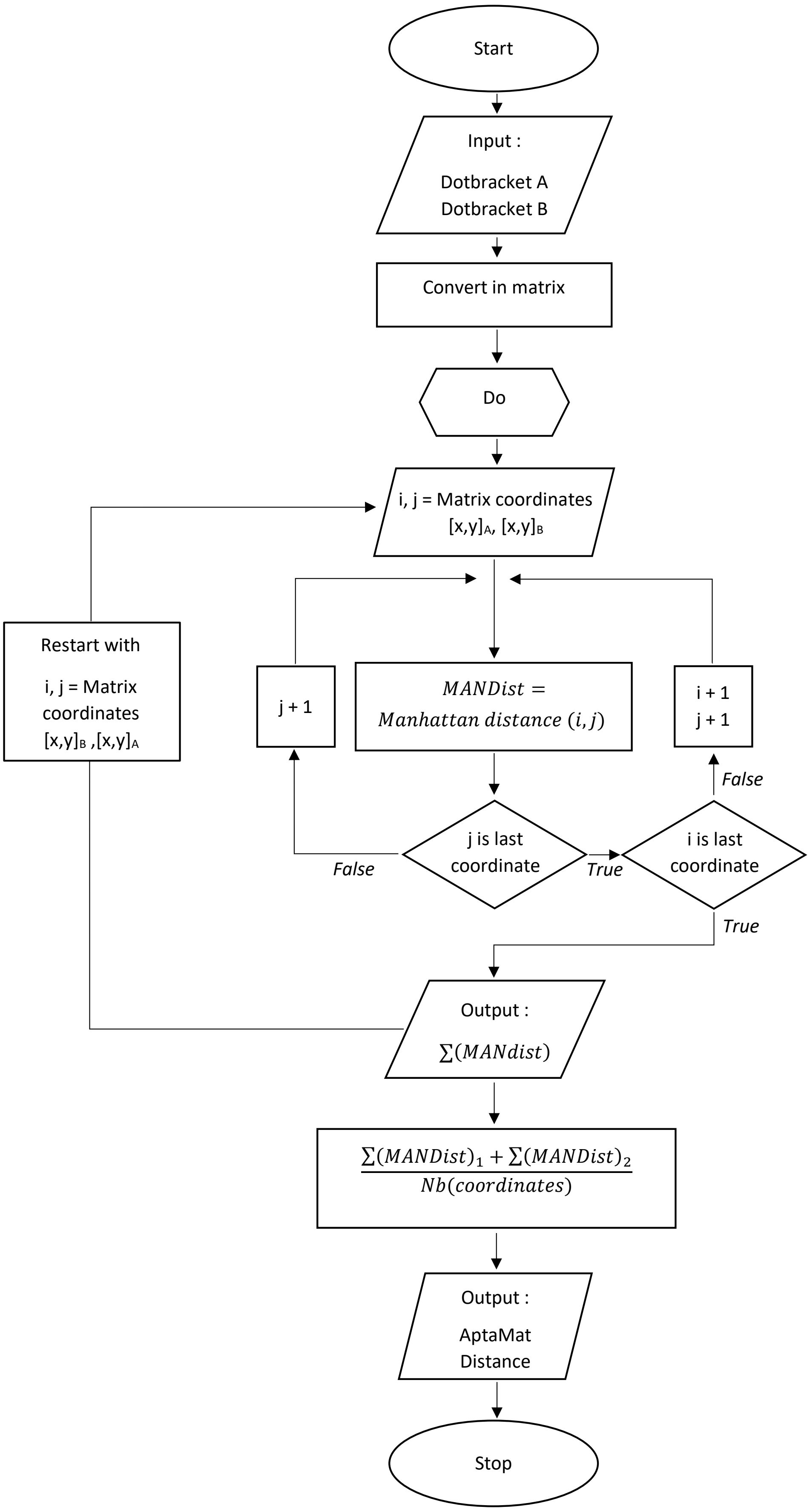


Figure S2: Flowchart of AptaMat algorithm

a)

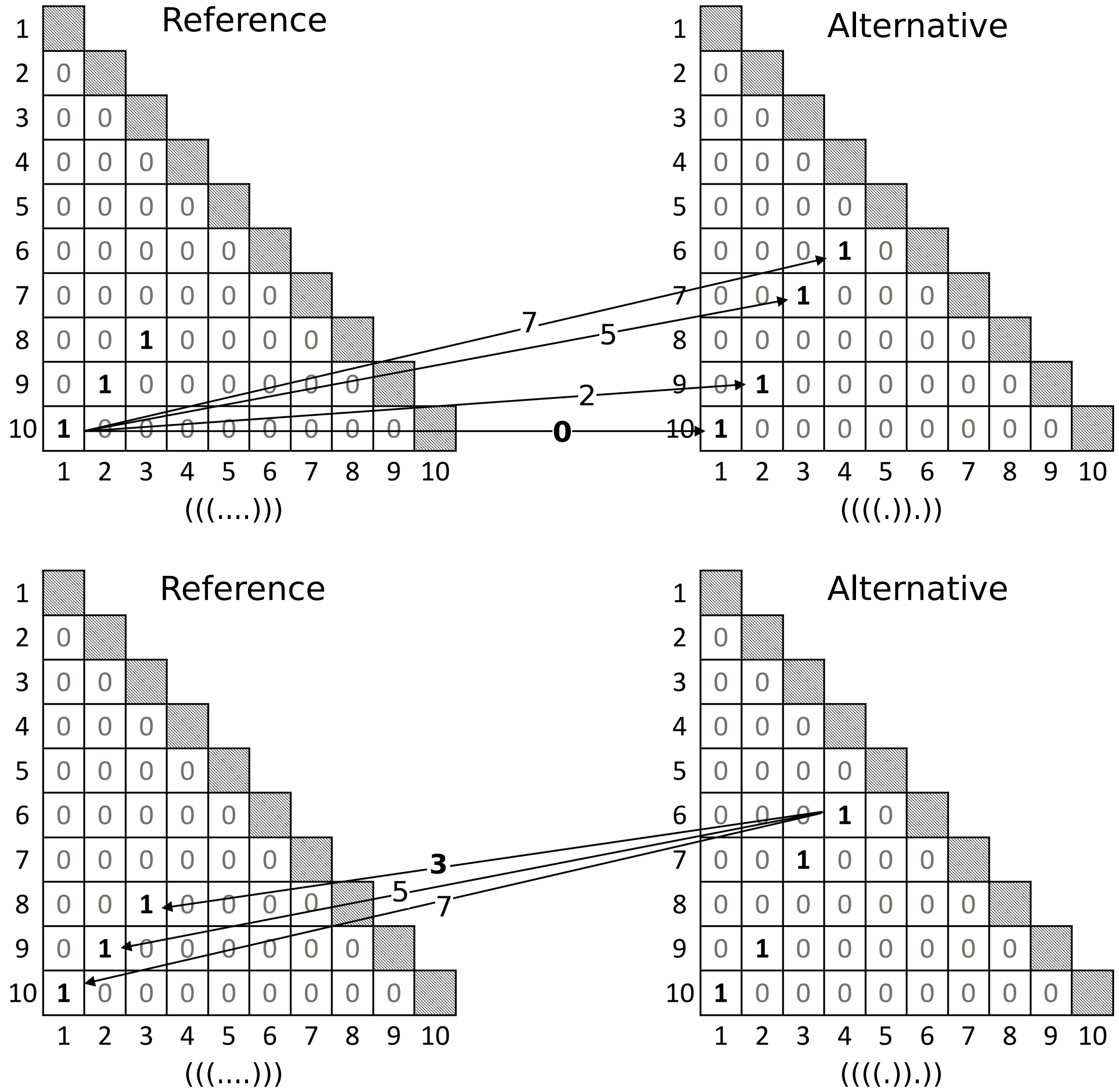


b)

| Coordinate | Reference: 1-10 | 2-9 | 3-8 | Lowest |
|---|---|---|---|---|
| *Alternative*: **1-10** | 0 | 2 | 4 | 0 |
| **2-9** | 2 | 0 | 2 | 0 |
| **3-7** | 5 | 3 | 1 | 1 |
| **4-6** | 7 | 5 | 3 | 3 |
| **Lowest** | 0 | 0 | 1 | |

c)

$$\frac{\sum Lowest\ ManDist\ (Ref,\ Alt) + \sum Lowest\ ManDist\ (Ref,\ Alt)}{nb\ coordinates}$$

$$= \frac{(\,0+0+1\,)+(\,0+0+1+3\,)}{7} = 0.714$$

Figure S3: Detailed AptaMat calculation for a simple example. a) Matrix representation of 2 secondary structures and the performed comparison shown by the arrows and their associated Manhattan distances. For each “1” from a matrix we find the nearest “1” from the second matrix in Manhattan distance, in both directions Reference –> Alternative, then Alternative –> Reference. b) The Manhattan distances between every pair of “1”s. c) The general formula to compute AptaMat and the resulting value for this example.

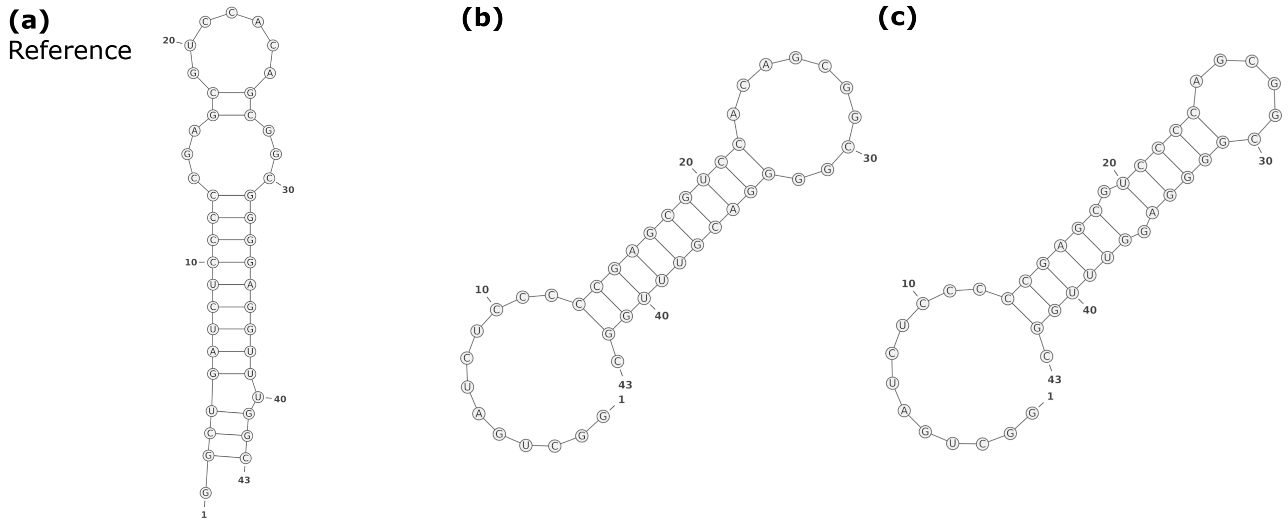


.((((((((((((...((.......))...))))))))).)))    ............((((((((((..........))))))))))).    ............((((((.(((((......)))))).))))).

| | | |
|---|---|---|
| Hamming distance | 0.56 | 0.56 |
| RNAdistance | 12 | 22 |
| DoPloCompare | 0.36 | 0.19 |
| AptaMat | 10.8 | 10.5 |

Figure S4: Graphical representations, dot-bracket notations of the representative structure of ssNA 3 (**a**) and the alternative structures (**b**) and (**c**). The Hamming distance, RNAdistance, DoPloCompare, and AptaMat distances from the reference are also reported.

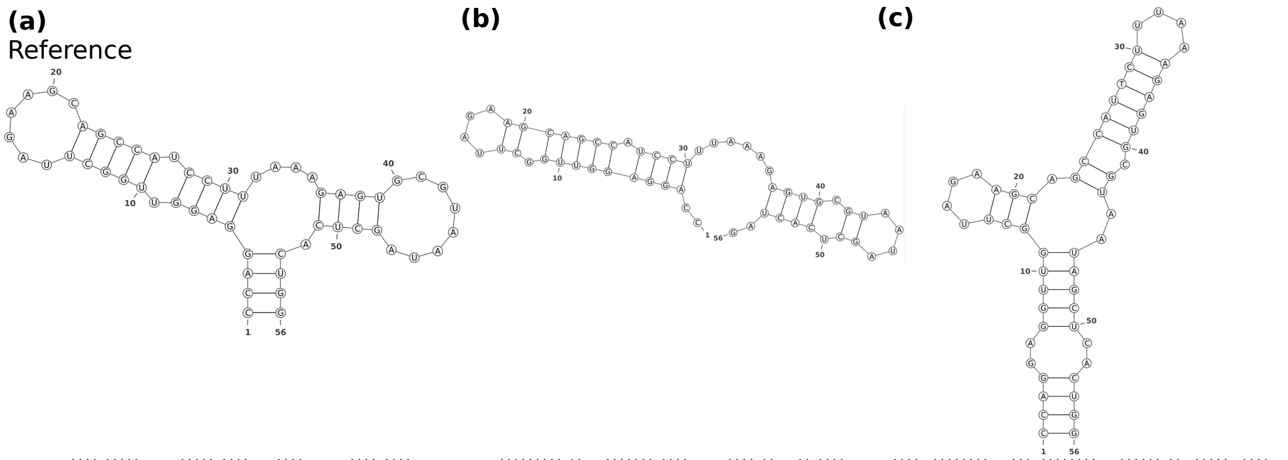


....((((.(((((.......))))).))))....((((........)))).))))    ..(((((((((.((....))))))).))))......((((.((....)).))))..    ((((..((((((((....))).((((((((....)))))).))..)))))..))))

| | | |
|---|---|---|
| Hamming distance | 0.32 | 0.48 |
| RNAdistance | 26 | 44 |
| DoPloCompare | 0.16 | 0.13 |
| AptaMat | 6.17 | 10.65 |

Figure S5: Graphical representations, dot-bracket notations of the representative structure of ssNA 4 (**a**), and the alternative structures (**b**) and (**c**). The Hamming distance, RNAdistance, DoPloCompare, and AptaMat distances from the reference are also reported.

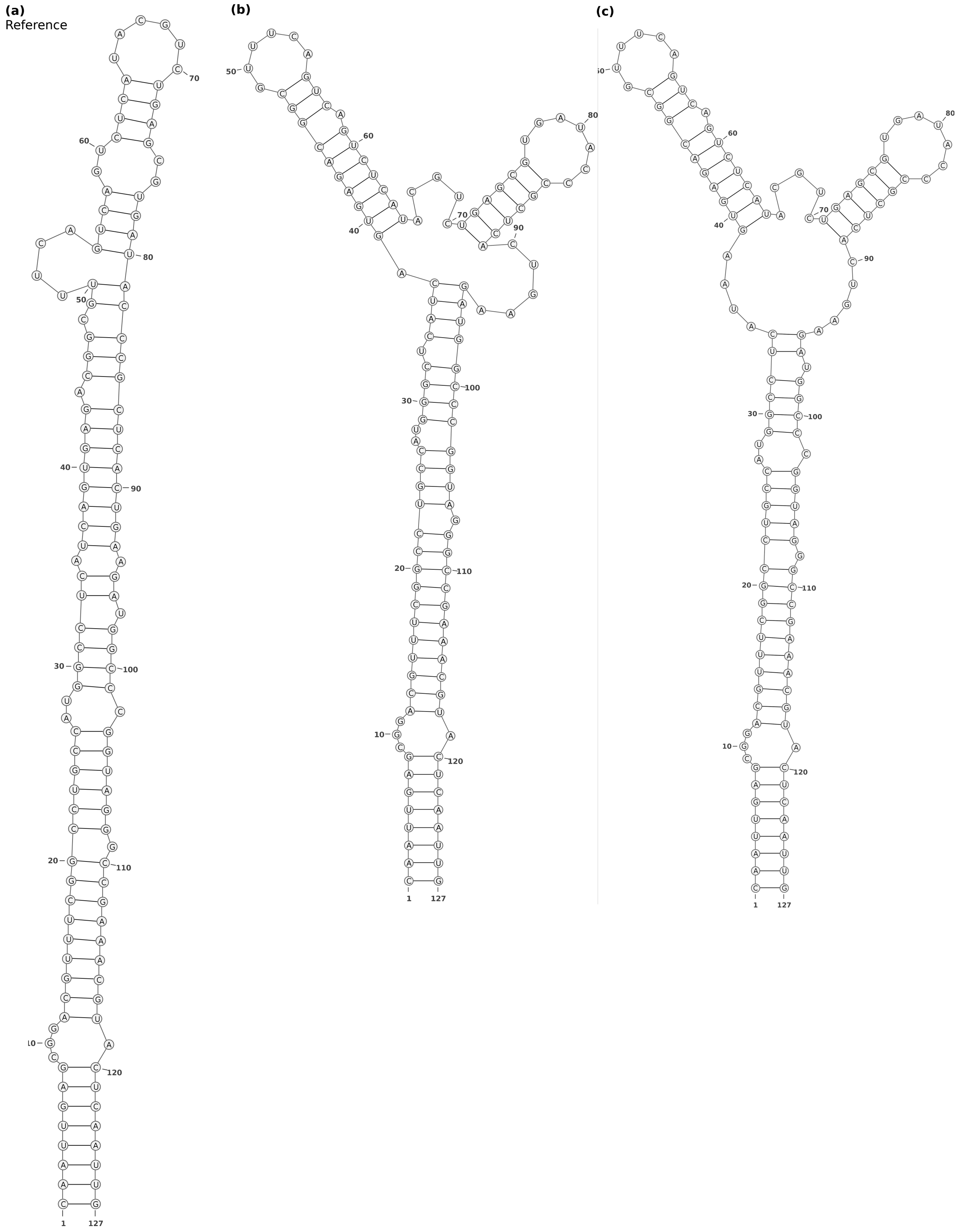


| | (a) | (b) | (c) |
|---|---|---|---|
| | ((((((((...((((((((((((((((..((((((.((((((((.(((.((....((((.. ((((......))))..)))))))))))))))))).)).)))).))))))).)))))))))).))))))))) | ((((((((...((((((((((((((((..((((.((((.((((((((((......))).))) ))))......((((((.......))))))......))))))))))))).))))))))))).)))))))) | ((((((((...((((((((((((((((..((((((....((((((((((......))).)))) )))......((((((.......))))))......)).)))).))))).))))))))))).)))))))) |
| Hamming distance | | 0.31 | 0.29 |
| RNAdistance | | 60 | 62 |
| DoPloCompare | | 0.023 | 0.022 |
| AptaMat | | 8.41 | 8.65 |

Figure S6: Graphical representations, dot-bracket notations of the representative structure of ssNA 5 (**a**), and the alternative structures (**b**) and (**c**). The Hamming distance, RNAdistance, DoPloCompare, and AptaMat distances from the reference are also reported.

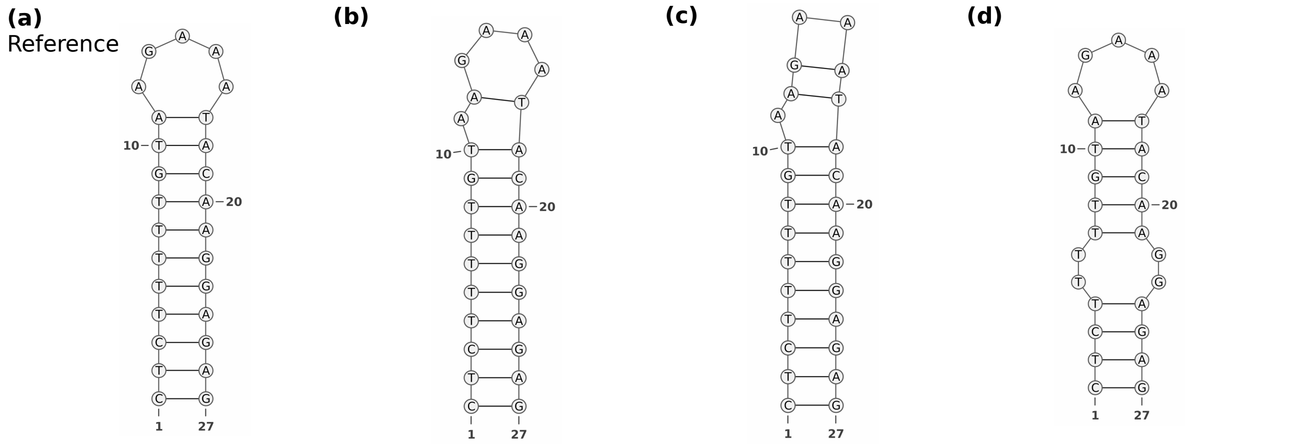

| | (a) Reference | (b) | (c) | (d) |
|---|---|---|---|---|
| | (((((((((((.....))))))))))) | ((((((((((.(....))))))))))) | ((((((((((.((..)))))))))))) | ((((..(((((.....)))))..)))) |
| Hamming distance | | 0.074 | 0.15 | 0.15 |
| RNAdistance | | 2 | 6 | 8 |
| AptaMat | | 0.091 | 0.22 | 0.20 |

Figure S7: SsNA 6 shows the limits of the Hamming distance and RNAdistance in comparing ssNAs secondary structures. Alternative structures **(c)** and **(d)** have the same Hamming distance to the reference secondary structure **(a)**, although structure **(d)** doesn't have the 5 nucleotides loop and it has a small bulge instead. The alternative structure **(c)** has a higher RNAdistance to the reference secondary structure **(a)**, although it only has 2 base pairs less than the reference structure. AptaMat, on the opposite, is able to correctly rank structures **c** and **d**.

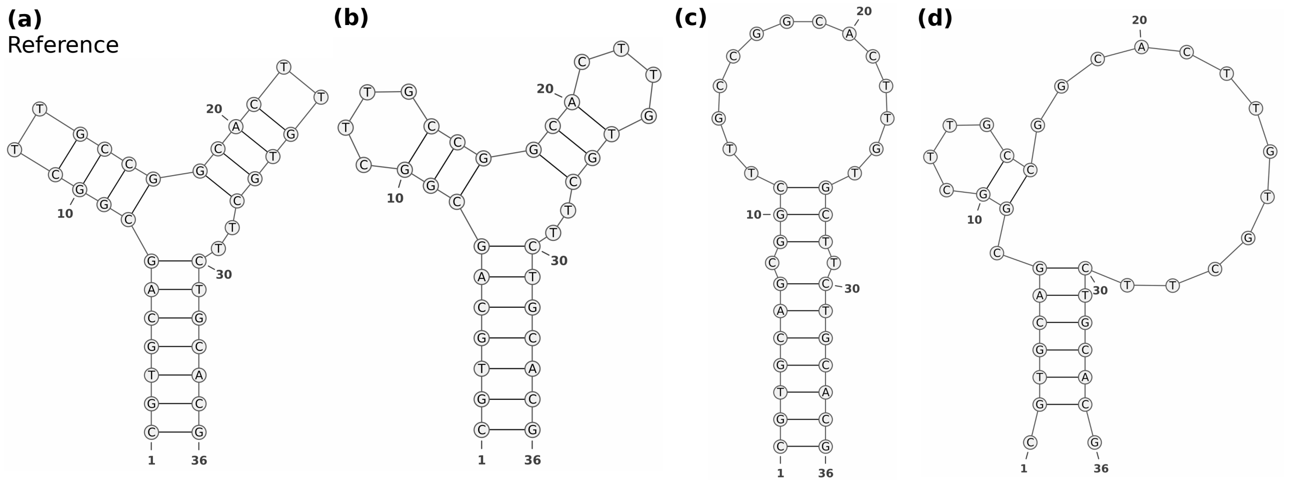

| | (a) Reference | (b) | (c) | (d) |
|---|---|---|---|---|
| | (((((((((((..))))((((..))))..))))))) | ((((((((((....)))(((....)))..))))))) | (((((((.(((..............))).))))))) | .((((((.((....)).............)))))). |
| Hamming distance | | 0.11 | 0.33 | 0.39 |
| RNAdistance | | 8 | 24 | 28 |
| AptaMat | | 0.14 | 4.16 | 3.21 |

Figure S8: Graphical representations, dot-bracket notations of the representative structure of ssNA 8 **(a)** and the alternative structures **(b)**, **(c)**, and **(d)**. The Hamming distance, RNAdistance, and AptaMat distances from the reference are also reported.

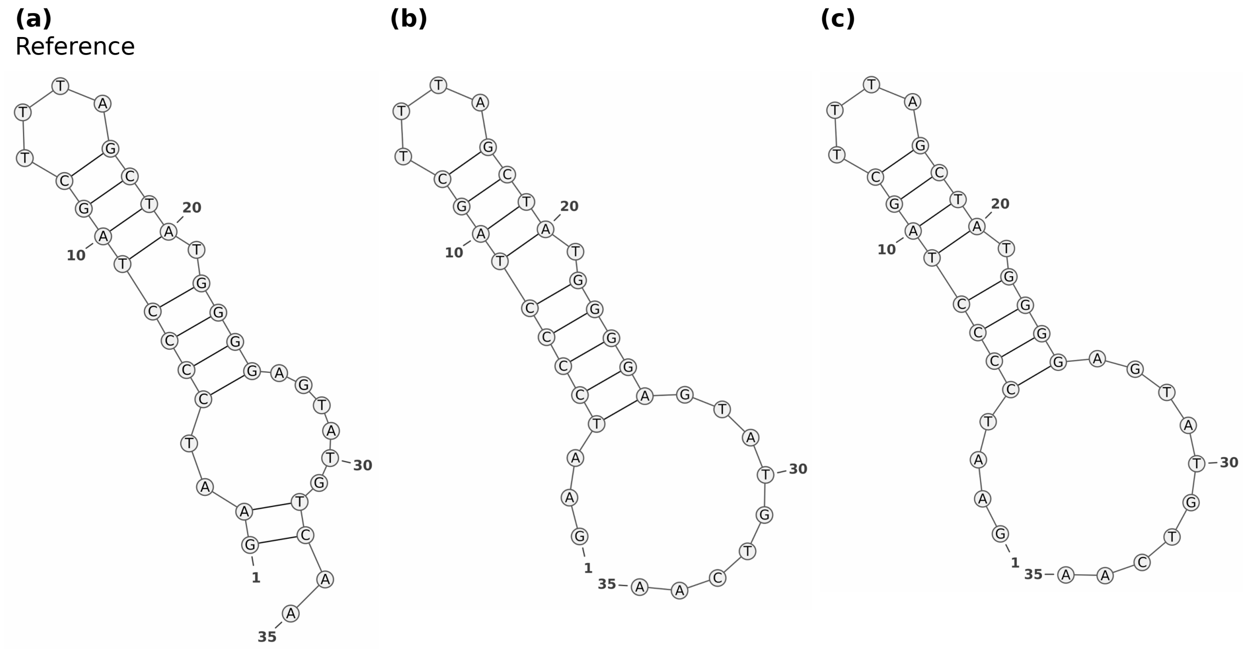


| | ((..((((((((....)))).))))......)).. | ...(((((((((....)))).)))))......... | ....((((((((....)))).))))......... |
|---|---|---|---|
| Hamming distance | | 0.17 | 0.11 |
| RNAdistance | | 8 | 8 |
| AptaMat | | 1.05 | 1.22 |

Figure S9: Graphical representations, dot-bracket notations of the representative structure of ssNA 9 (**a**) and the alternative structures (**b**) and (**c**). The Hamming distance, RNAdistance, and AptaMat distances from the reference are also reported.

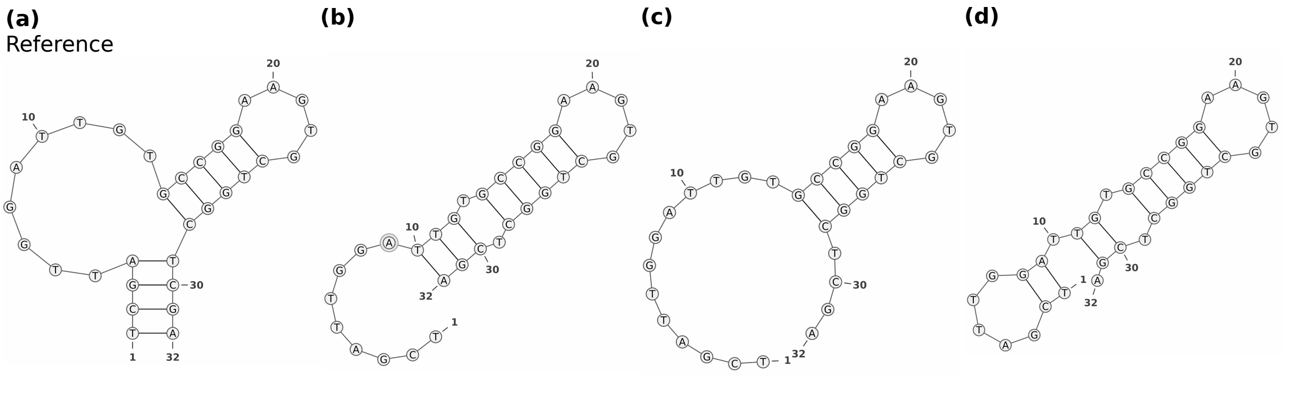


| | ((((.........(((((.....))))))))) | .........(((.(((((.....))))).))) | .............(((((.....))))).... | ((.....)).((.(((((.....))))).)). |
|---|---|---|---|---|
| Hamming distance | | 0.25 | 0.25 | 0.25 |
| RNAdistance | | 16 | 16 | 20 |
| AptaMat | | 3.18 | 4 | 5.22 |

Figure S10: Graphical representations, dot-bracket notations of the representative structure of ssNA 10 (**a**) and the alternative structures (**b**), (**c**), (**d**), and (**e**). The Hamming distance, RNAdistance, and AptaMat distances from the reference are also reported.

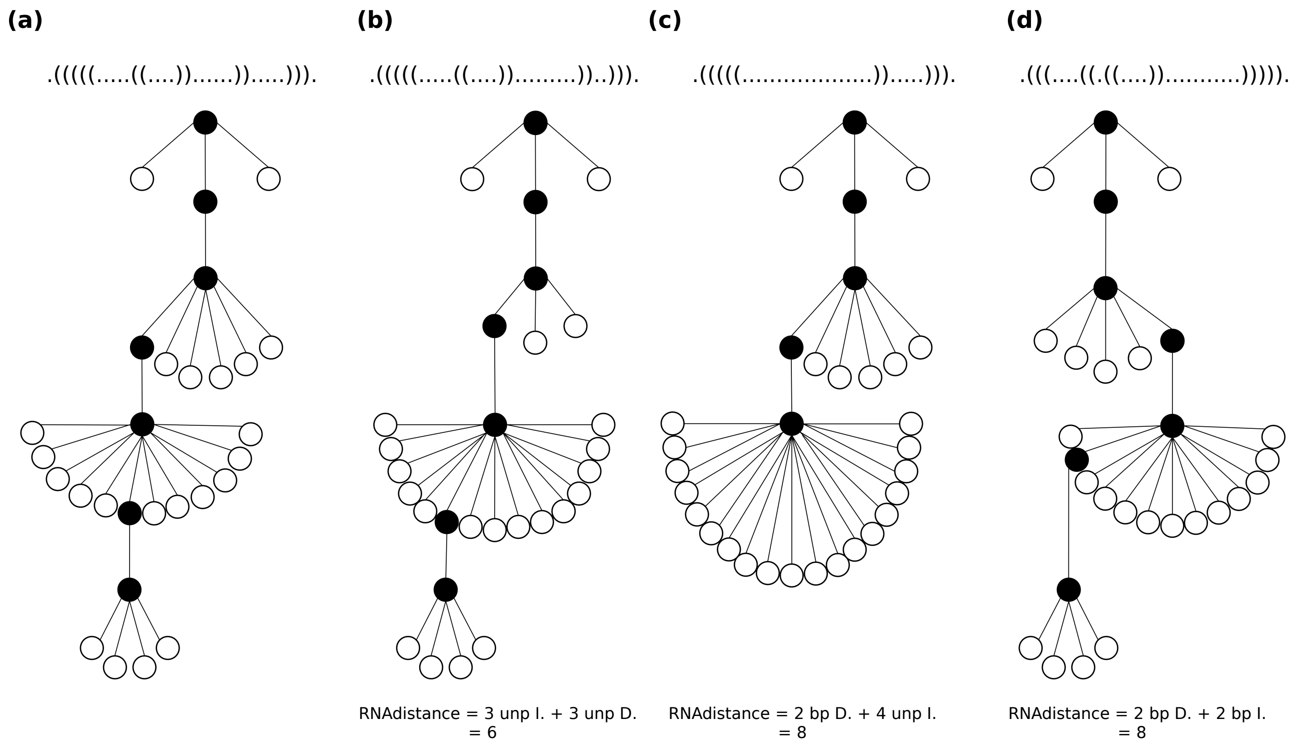


Figure S11: Dot-bracket and tree representations of ssNA 11 (**a**) and the alternative structures (**b**), (**c**), and (**d**). The RNAdistance calculation are also reported, where "bp" means base pair, "unp" means unpaired base, "I." means insertion, and "D." deletion.

#### References

[Ivry *et al.*, 2009] Ivry, T., and Michal,S., Avihoo,A., Sapiro, G., Barash, D. (2009) An image processing approach to computing distances between RNA secondary structures dot plots., *Algorithms for Molecular Biology*, **4**, 4.
